# Magnetic particle imaging reveals heterogeneous retention, leakage and redistribution of nanoparticles following intratumoral injection

**DOI:** 10.64898/2026.05.13.724909

**Authors:** Ali Shakeri-Zadeh, Asif Itoo, Janani Gurumurthy, Preethi Korangath, Robert Ivkov, Jeff W.M. Bulte

**Author notes:** These authors contributed equally (Ali Shakeri-Zadeh, Asif Itoo).

## Abstract

Intratumoral (i.t.) delivery of nanoparticles (NPs) is widely used to achieve high local NP concentrations. However, the temporal fate of i.t.-injected NPs remains poorly understood. We present a quantitative approach using whole-body magnetic particle imaging (MPI) to track magnetic NPs (MNPs) following i.t. injection. Using fiducial-calibrated imaging, we quantified MNP mass over time in subcutaneous 4T1 breast tumors. Longitudinal imaging revealed progressive loss of i.t. MNP content and heterogeneous systemic redistribution across animals despite standardized delivery conditions. *Ex vivo* MPI confirmed off-target accumulation primarily in the liver and spleen, consistent with reticuloendothelial clearance pathways. Histological analysis demonstrated spatially heterogeneous i.t. MNP deposition, potentially associated with local vascular features and tumor microenvironmental heterogeneity that may influence i.t. MNP retention or MNP clearance from the tumor. These findings highlight the importance of quantitative longitudinal whole-body MPI for understanding the fate of MNPs for informing localized nanotherapy.

## INTRODUCTION

Intratumoral (i.t.) injection is an established clinical strategy for achieving high local drug concentrations in solid tumors while minimizing systemic exposure. I.t. delivery addresses key limitations of systemic delivery including vascular barriers, rapid clearance, and off-target biodistribution.^1, 2^ This delivery method has been clinically explored across multiple solid tumor types^3^ using diverse therapeutic modalities, including viral immunotherapy,^4^ cytokines,^5^ checkpoint inhibitors,^6^ radiotherapeutics,^7^ and magnetic fluid hyperthermia (MFH),^8^ underscoring its expanding role in localized cancer therapy. Among these approaches, MFH represents a uniquely distribution-dependent modality, as therapeutic heating scales directly with the spatial concentration and total mass of magnetic nanoparticles (MNPs). MNPs have been directly injected into glioblastoma^8^ and prostate^9^ tumors in clinical trials, demonstrating the feasibility, safety, and therapeutic potential of i.t. MNP delivery for localized MFH.

Despite these advances, most i.t. delivery strategies implicitly assume that injected agents remain spatially confined within the tumor volume, an assumption that has not been rigorously validated *in vivo*.^10-13^ Following i.t. injection, NPs may exhibit highly heterogeneous i.t. distribution^14, 15^ governed by injection location,^16^ tumor vascularization, extracellular matrix density, and elevated interstitial fluid pressure.^17^ These factors can promote leakage into systemic circulation or surrounding tissues, thereby reducing effective i.t. dose delivery and introducing uncertainty in both therapeutic efficacy and safety. For example, irregular vasculature^18^ and elevated interstitial pressure^19^ in breast tumors can promote escape of i.t. injected agents from the injection site into surrounding drainage pathways or circulation. This redistribution may subsequently contribute to off-target uptake in organs such as the liver and spleen, thereby compromising the intended localized therapeutic effect.

Noninvasive longitudinal monitoring of i.t. retention and systemic leakage is therefore equally essential for i.t. delivery as pharmacokinetic evaluation is for systemic injection.^14, 20-22^ Measurable redistribution to off-target organs following i.t. injection has been reported for different NP platforms including radiolabeled nanoshells,^23^ lipid NPs,^11^ and MNPs.^15, 24^ However, these studies have relied on endpoint measurements and have not resolved the longitudinal whole-body kinetics or inter-tumoral variability of NP leakage and *in vivo* redistribution. Moreover, current clinical imaging modalities lack the sensitivity, quantitative accuracy, or repeatable whole-body capability required for this task.^20, 22^ Magnetic particle imaging (MPI) provides background-free signal generation with linear quantification of MNP mass, enabling sensitive and repeatable *in vivo* tracking of their biodistribution over extended time periods.^21, 25-27^ These properties uniquely position MPI as a quantitative tool for directly measuring NP retention and redistribution dynamics following i.t. delivery.^14, 25^

Here, we present a longitudinal MPI-based framework to quantitatively track the retention and leakage dynamics of i.t.-delivered MNPs in subcutaneous 4T1 breast tumors. Using fiducial-based calibration combined with whole-body MPI, we aimed to achieve accurate quantification of i.t. iron content and its redistribution over time. Longitudinal measurements over eight days revealed progressive MNP clearance from the tumor with high inter-subject variability as a result of underlying tumor heterogeneity. This work provides quantitative insight into the transport kinetics of locally delivered NPs and presents a platform applicable to optimizing dosing strategies, safety profiles, and translational design of i.t. nanotherapies.

### EXPERIMENTAL

#### MPI calibration and quantitative signal normalization

Quantitative MPI measurements were performed using a Momentum™ MPI scanner (Magnetic Insight Inc., Alameda, CA, USA). A fiducial-based calibration approach was used to convert MPI signal into iron mass. The stock of Synomag®-D70 NPs (micromod Partikeltechnologie GmbH, Germany) with a reported concentration of 12 mg Fe/mL was re-analyzed using a Ferrozine-based spectrophotometric chemical assay.^14, 26^ Four fiducial samples containing known iron masses (30, 60, 120, and 240 µg Fe in 20 µL PBS) were prepared and positioned in a 3D-printed custom holder^28^ to maintain fixed placement within the MPI field of view during imaging. MPI scans were acquired using standard 2D acquisition mode. 2D imaging was selected because our prior work^28^ demonstrated this acquisition strategy minimized “spillover effects” and provided more robust quantitative measurements for longitudinal MNP tracking compared to other MPI scan modes on our instrument. Regions of interest (ROIs) were delineated around fiducial signals, and total MPI signal was calculated as mean intensity × ROI area. A calibration curve was generated by linear regression of MPI signal versus iron mass and used for all subsequent conversion of MPI measurements to iron content.

#### 4T1 cell culture

Murine 4T1 mammary carcinoma cells were cultured under standard conditions as previously described.^15^ Cells were maintained in RPMI-1640 medium supplemented with 10% fetal bovine serum and 1% penicillin–streptomycin at 37 °C in a humidified incubator with 5% CO_2_. Cells were passaged at sub-confluent densities using trypsin-EDTA detachment and routinely inspected to ensure normal morphology and growth characteristics.

#### Animal model, tumor implantation, and i.t. injection

All procedures were performed in accordance with institutional guidelines and were approved by our University’s Animal Care and Use Committee. Female BALB/c mice (6–8 weeks old) were used to establish syngeneic 4T1 breast tumors as previously described.^15^ Briefly, 4T1 cells were harvested during logarithmic growth phase and were resuspended in sterile PBS and injected subcutaneously into the flank (n = 10 mice). Tumors were allowed to grow to 200-250 mm^3^ prior to i.t. injection and longitudinal imaging. Tumor dimensions were measured using a digital caliper (measurement uncertainty: ±0.02 mm), and tumor volume was calculated using the standard ellipsoid approximation formula: V=(π/6)×L×W×H. I.t. injection of MNPs was performed using the same batch of Synomag®-D70 that was characterized during the calibration step to ensure consistency between calibration and *in vivo* measurements. At the time of i.t. injection and initiation of MPI studies, four mice with comparable tumor volumes were selected (231.2 mm^3^ for M1; 230.3 mm^3^ for M2; 231.9 mm^3^ for M3; and 215.7 mm^3^ for M4). I.t. MNP injection was performed in all four mice. Each selected tumor received a single i.t. injection of 10 µL suspension corresponding to a total dose of 120 µg Fe. Injections were performed using a Hamilton syringe connected to a syringe pump (Pump 11 Elite, Harvard Apparatus, Holliston, MA, USA) set at a controlled infusion rate of 1 µL/min^15^ to minimize backflow due to interstitial pressure. The needle was inserted approximately into the central region of the tumor with care taken to promote localized deposition and reduce visible reflux during needle withdrawal. Following injection, animals were immediately prepared for MPI at the first time point (15 min post-injection). However, data from M4 were excluded from subsequent analysis because the measured MPI-derived tumor iron content exceeded the injected dose by approximately 78%. The source of this discrepancy remains unclear and warrants further investigation in well-designed phantom and *in vivo* studies. Such overestimation may reflect MPI quantification artifacts associated with closely spaced high-signal regions in the tumor, causing sub-centimeter spillover effects that we previously reported for nearby MPI hotspots (distances < 2 cm for standard MPI scans).^28^

#### Longitudinal whole-body MPI

Imaging was initiated 15 min following i.t. injection and repeated at 3 h, 6 h, and 1, 2, 4, and 8 days post-injection to monitor MNP retention and redistribution over time. Serial imaging was conducted using standard 2D MPI acquisition mode to enable longitudinal tracking while minimizing scan duration and maintaining consistent imaging conditions across sessions.^28^ Animals were anesthetized with inhaled isoflurane (1–2% in medical air/oxygen) during imaging and positioned in a dedicated holder^28^ to ensure reproducible placement within the field of view. The same four fiducial references used for calibration were included in every imaging session and maintained at fixed positions relative to the imaging field. At the terminal imaging time point (day 8 post-injection), additional whole-body 3D MPI scans (21 radial slices in standard mode) were acquired to assess spatial distribution of MNPs and confirm organ-level redistribution observed during longitudinal 2D imaging. Following MPI acquisition, anatomical computed tomography (CT) images were collected using an IVIS Spectrum/CT imaging system (Caliper Sciences, Waltham, MA, USA). Imaging parameters were kept consistent across all MPI sessions.

#### Image processing and longitudinal quantification

Two-dimensional MPI datasets were analyzed using ImageJ (National Institutes of Health, Bethesda, MD, USA). For each imaging time point, ROIs were manually delineated around MPI signal hot spots corresponding to tumor regions, based on the high signal-to-background inherent to MPI. When present, additional ROIs were drawn over anatomically consistent regions exhibiting off-target signal to assess MNP redistribution. Total MPI signal was quantified as the product of mean signal intensity within each ROI and the corresponding ROI area. Iron content was obtained by converting MPI signal values using the fiducial-based calibration curve described above. Measurement uncertainty was estimated from spatial variability of signal intensity within each ROI. Longitudinal tumor retention curves were generated by plotting calculated iron mass and MPI signal intensity over time using Microsoft Excel (Microsoft Corp., WA, USA). 3D MPI datasets were co-registered with corresponding micro-CT images using 3D Slicer software (version 5.10.0; www.slicer.org) as previously described.^14, 26^

#### *Ex vivo* MPI

At the terminal imaging time point, animals were euthanized and tumors together with organs exhibiting *in vivo* MPI signal (liver and spleen) were excised for *ex vivo* analysis. Tissues were gently rinsed in PBS to remove residual blood and individually scanned by *ex vivo* MPI. Imaging was conducted using standard acquisition settings consistent with *in vivo* scans to allow direct comparison. ROIs were manually delineated around each tissue sample using ImageJ, and total MPI signal was quantified as described above. Iron content was calculated using the fiducial-based calibration curve to assess MNP retention and redistribution.

#### Prussian blue (PB) and anti-CD31 staining

After *ex vivo* imaging, tissues were immersed in 4% PFA at 4 °C for 24 h, and then transferred to 30% sucrose for another 72 h. After embedding in optimal cutting temperature compound, tissues were sequentially cut into 20 μm sections using a cryostat (Thermo Fisher Scientific). For PB staining, tissues were fixed with 4% glutaraldehyde for 20 min, washed with PBS, and incubated with Perl’s reagent (potassium ferrocyanide in HCl) for 30 min at room temperature.^14, 26^ To enhance tissue visualization, sections were counterstained with nuclear fast red (NFR) for 20 mins. Stained sections were washed with deionized water to remove excess reagent and then imaged using a Zeiss Apotome 2 microscope. Quantitative analysis was performed by threshold-based pixel segmentation using ImageJ to determine total tissue area and PB-positive area, allowing estimation of iron coverage within tumor and organ sections.

To evaluate tumor vascular architecture, sequential tissue sections were subjected to immunofluorescence staining for CD31 (platelet endothelial cell adhesion molecule-1) as a vascular endothelial marker. Tissue sections were first blocked with 3% bovine serum albumin (BSA) in Tris-buffered saline (TBS, pH = 7.0) for 1 h at room temperature to reduce nonspecific binding. Sections were then incubated with rat anti-mouse CD31 primary antibody (1:40 dilution; Dianova, DIA 310) prepared in 3% BSA in TBS overnight at 4 ºC. Sections were washed three times with 1× TBS and subsequently incubated with goat anti-rat Alexa Fluor 594 secondary antibody (1:500 dilution; Invitrogen, A11007) prepared in 3% BSA in TBS for 1 h at room temperature. Sections were washed three times with 1× TBS and mounted with an antifade mounting medium containing DAPI to counterstain nuclei. Fluorescence images were acquired using a Zeiss Axiovert 200 M inverted epifluorescence microscope equipped with an Axiocam camera under identical imaging settings across samples. Quantitative analysis of vascular density was performed using ImageJ software by threshold-based segmentation of CD31-positive pixels relative to the total tissue area.

## RESULTS AND DISCUSSION

### Quantitative calibration enables MPI-based MNP measurement

Reliable longitudinal quantification of i.t. MNP retention and systemic redistribution requires a direct and reproducible relationship between MPI signal and iron mass. Fiducial-based calibration demonstrated a strong linear dependence of MPI signal on iron content across the experimentally relevant range (**Fig. 1**), confirming that MPI signal intensity can be used as a quantitative surrogate for iron mass.^27, 29, 30^ Importantly, the injected iron dose used for *in vivo* studies fell within the calibrated dynamic range, ensuring that subsequent measurements were derived from interpolation rather than extrapolation. Maintaining identical fiducial references throughout the study enabled session-to-session normalization, ensuring that longitudinal signal changes primarily reflected i.t. MNP retention and systemic redistribution rather than instrumentation variability. Accordingly, the same four fiducials were included in all subsequent *in vivo* imaging sessions.

**Figure 1:**
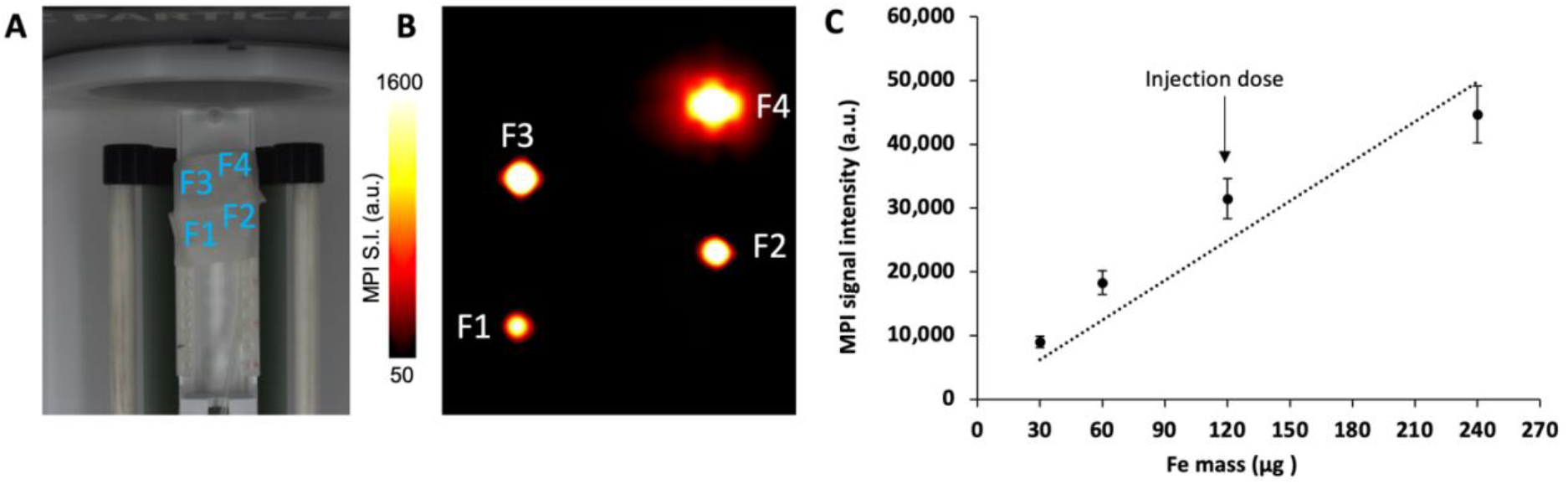
Calibration setup for quantitative MPI signal normalization. (**A**) Experimental setup showing the four fiducials (F) secured within the custom holder (asterisks). (**B**) Representative 2D MPI image of fiducials containing known iron mass for quantitative calibration. (**C**) Linear calibration curve correlating MPI signal intensity (defined as mean signal intensity within the ROI multiplied by ROI area) with iron mass, enabling conversion of MPI signal to iron content (y = 207.46×, R^2^ = 0.967). The calibration range is within the range of the injected *in vivo* iron dose to ensure quantitative accuracy. Error bars represent the standard deviation of MPI signal intensity within each ROI propagated through the signal × area calculation

### Time-dependent i.t. MNP retention and redistribution following i.t. injection

Longitudinal MPI demonstrated that i.t. MNP retention evolved progressively rather than remaining spatially confined (**Fig. 2**). Across all animals, tumor-associated signal decreased steadily over time, indicating redistribution of MNP from the injection site, with mean tumor iron retention declining by approximately 1.8-to 3.0-fold between 15 min and day 8. This temporal behavior is consistent with post-injection MNP redistribution processes, likely reflecting early pressure-driven dispersion followed by diffusive and convective redistribution through interstitial pathways.^17^ Despite nearly identical tumor volumes (230-232 mm^3^) and standardized injection conditions, longitudinal trajectories differed among the three animals, suggesting the influence of tumor heterogeneity on i.t. MNP retention.^16-18^

**Figure 2:**
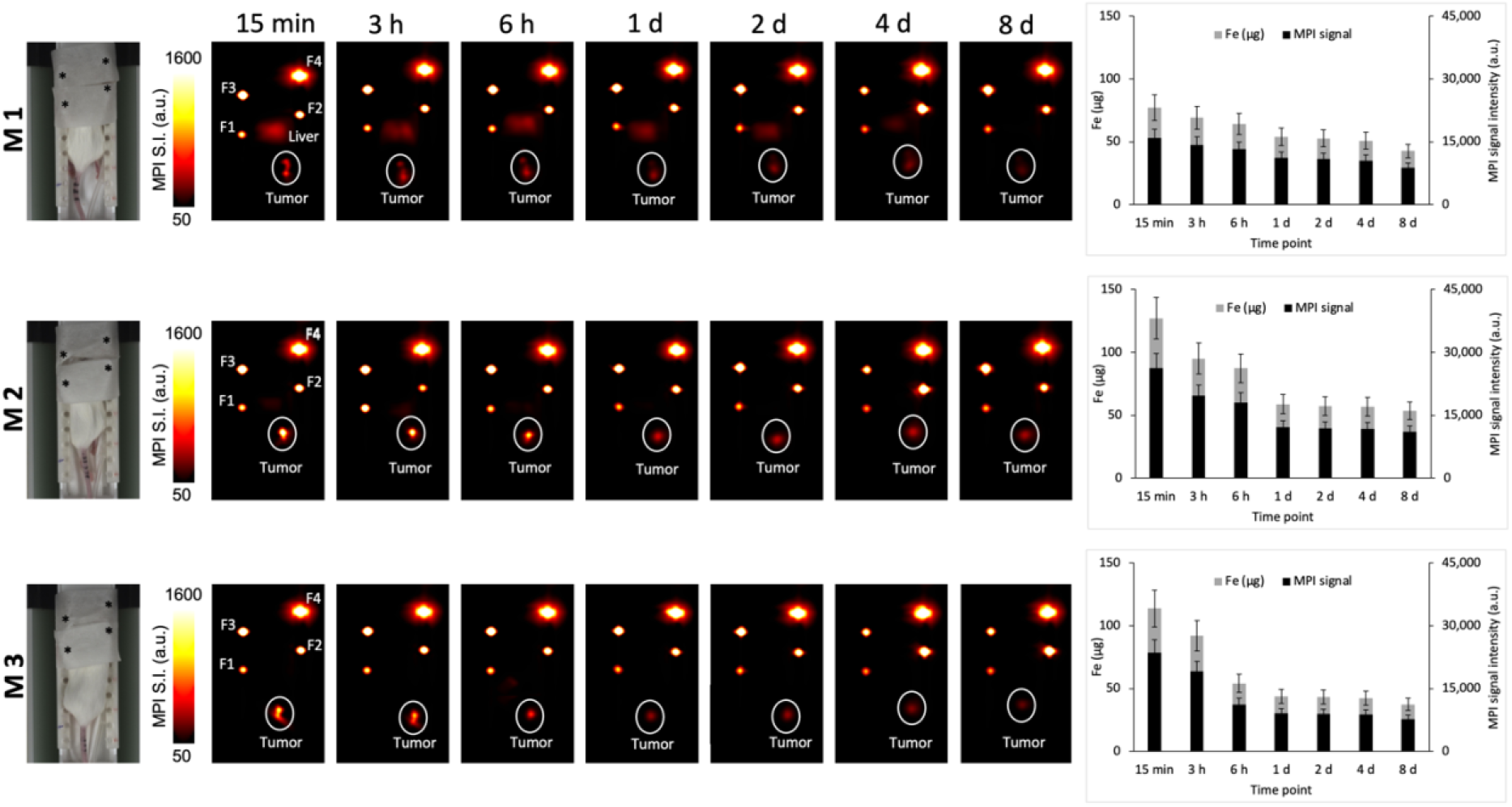
Longitudinal whole-body MPI of i.t. MNP retention and systemic redistribution. Serial MPI scans are shown for three 4T1 tumor-bearing mice at multiple time points following i.t. injection of Synomag®-D70 (10 µL, 120 µg Fe). The same fiducial references (F1–F4) were used for quantitative normalization across imaging sessions. The tumor region is outlined in white. Right panels show longitudinal quantification of MPI signal intensity and corresponding calculated iron mass within tumor regions derived from fiducial-calibrated measurements. Error bars represent the standard deviation of MPI signal intensity within each tumor ROI propagated through the signal × area calculation.

Our findings shown in **Fig. 2** also underscore a critical limitation of conventional i.t. dosing paradigms: the injected dose alone does not determine the design of nanotherapy parameters over time. Redistribution significantly alters the retained local MNP mass and therefore affects the setup of therapeutic settings of different nanotherapy methods, including MFH,^31^ NP-mediated radiosensitization,^32^ and NP-based drug delivery.^11^ Longitudinal quantification using whole-body MPI thus provides essential insight into treatment-relevant NP kinetics that cannot be inferred from static measurements.

In contrast to prior magnetic sensing studies,^12, 13^ which primarily quantified MNPs in tumors or individual organs separately and often at limited time points, our results demonstrate longitudinal whole-body MNP redistribution with substantially more dynamic and heterogeneous behavior following i.t. MNP delivery. Kettering et al.^13^ reported minimal redistribution of i.t. injected MNPs (fluidMAG-D, Fe_3_O_4_, starch-coated, ∼200 nm hydrodynamic diameter) over time using magnetorelaxometry, concluding that most MNPs remained confined within tumors with only limited accumulation in the liver and spleen and negligible systemic loss (less than 1% of injected NPs). While differences in MNP formulation and physicochemical properties may contribute to distinct biodistribution behavior, magnetorelaxometry provides bulk magnetic signal measurements with limited spatial localization, restricting its ability to resolve heterogeneous redistribution of MNPs within and beyond the tumor over time. The MPI-hyperthermia study by Kuboyabu et al.^12^ primarily evaluated therapy-associated signal changes in tumor and correlations with treatment response. While their work demonstrated the feasibility of using MPI to quantify i.t. signal and predict hyperthermia outcomes, it did not address whether MNPs remained spatially confined or how their distribution evolved after delivery.

### *Ex vivo* MPI confirms heterogeneous organ-level redistribution

To verify that longitudinal changes reflected true biological redistribution rather than methodological variability, *ex vivo* MPI was performed at the terminal *in vivo* imaging time point (**Fig. 3**).

**Figure 3:**
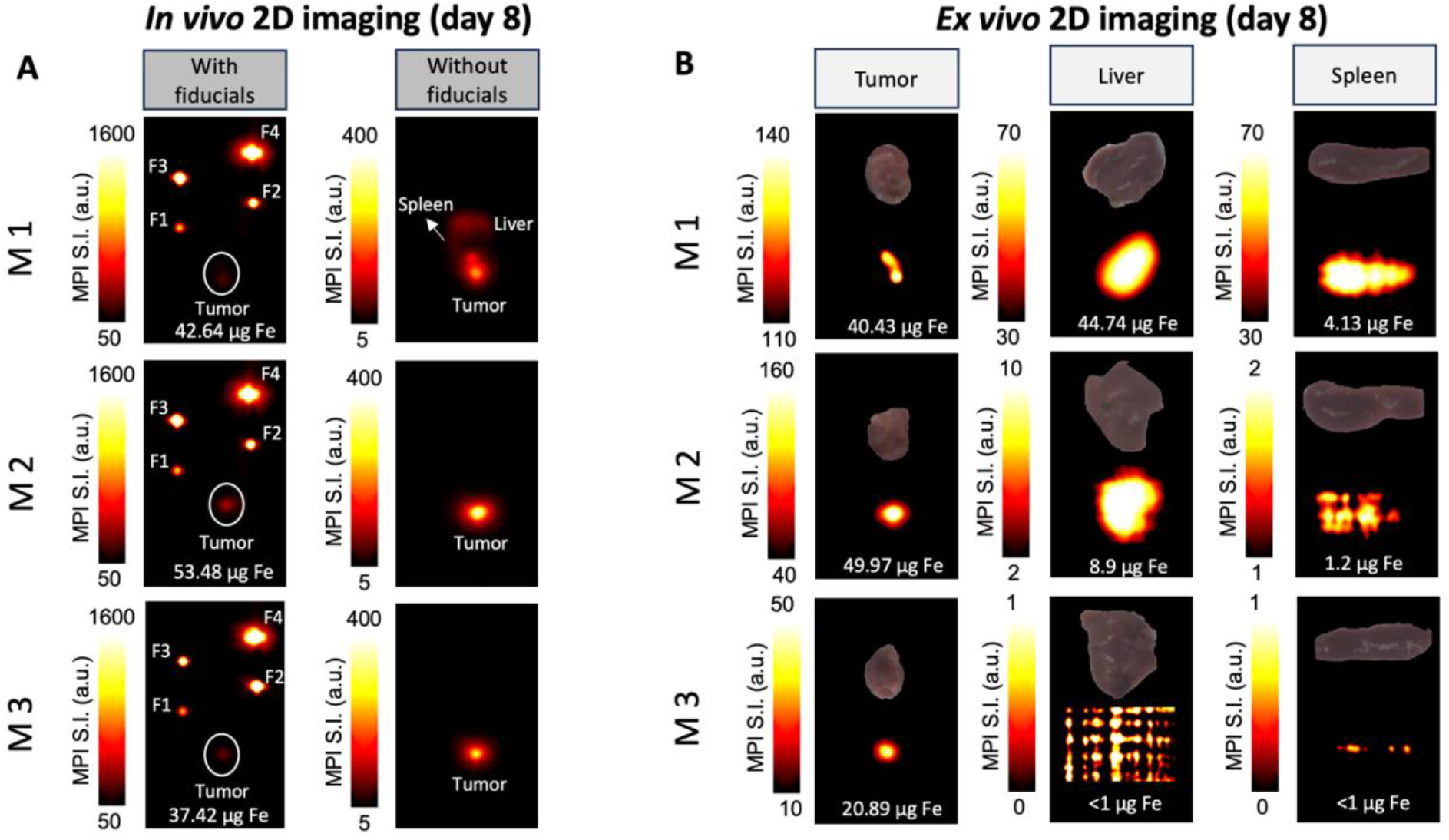
*In vivo* and *ex vivo* MPI of MNP retention and systemic redistribution eight days post i.t. injection. (**A**) *In vivo* whole-body MPI acquired at the terminal time point with and without fiducial references. Tumor regions are outlined with the corresponding quantified iron content. Imaging without fiducials with high MNP concentrations highlights off-target MNP redistribution, including detectable signal in the liver and spleen in M1. (**B**) *Ex vivo* MPI of the excised tumor, liver, and spleen from the same mice and quantification of retained MNPs. Measured iron content is shown for each tissue. Note, the lack of signal from liver and spleen in M3. *Ex vivo* imaging confirms heterogeneous MNP redistribution observed *in vivo*, with substantial inter-subject variability in tumor retention and systemic uptake. Color scales represent MPI signal intensity (a.u.) and were adjusted independently for visualization.

Organ measurements revealed inter-subject differences in MNP distribution patterns that were consistent with the *in vivo* MPI observations. Whole-FOV *in vivo* MPI signal (including mouse and fiducials) varied by less than ±7% across animals, supporting comparable overall detected MNP signal and suggesting that the observed differences primarily reflect heterogeneous MNP distribution rather than major procedural variability. Tumors retained measurable MNP fractions in all cases, but the extent of systemic accumulation varied markedly, with detectable liver and spleen signal in M1 and M2, and minimal (undetectable) off-target uptake in M3 at the terminal time point. *Ex vivo* MPI confirmed that redistribution occurred primarily into the reticuloendothelial system (RES), consistent with known clearance pathways for MNPs; however, other clearance/redistribution pathways/organs (e.g. bone marrow and lymph nodes) were not examined ex vivo and therefore cannot be excluded.^33-35^ Previous studies^34, 35^ have attributed hepatic accumulation of circulating MNPs primarily to sequestration by Kupffer cells, whereas splenic uptake has been associated with filtration and retention by red pulp macrophages within the splenic RES.

Although the present study includes a limited number of animals, this does not diminish the mechanistic relevance of the findings. Solid tumors are intrinsically heterogeneous systems with inter-individual variability, and strictly reproducible retention kinetics among individuals in larger cohorts are not expected even under identical delivery conditions, as tumor microenvironmental variability has been shown to significantly influence NP delivery and disposition across tumor models.^18^ The consistent decline in tumor MPI signal across animals supports the central finding, that even with consistent i.t. delivery, variable NP distributions can be expected. Variability in retention magnitude and systemic redistribution reflects physically and biologically meaningful differences in tumor-specific transport.

### A potential association between systemic redistribution and tumor vascular heterogeneity

Histological analysis (**Fig. 4**) was performed on two representative cases intentionally selected based on distinct *in vivo* MPI profiles (**Figs. 3** and **4**): M1 (tumor + RES signal) and M3 (tumor-only signal). These analyses were intended to provide illustrative spatial insight into MNP distribution patterns rather than statistical assessment across the cohort. Here, we acknowledge that histological findings from selected tissue sections cannot be directly compared with *in vivo* or *ex vivo* MPI measurements, as MPI reflects MNP distribution across the entire animal body or whole tumor volume, whereas histology provides localized information from a limited sample size that may not represent only a portion of the tumor distribution. In our study, PB staining demonstrated spatially heterogeneous MNP deposition within the tumor sections, suggesting localized iron accumulation patterns following i.t. injection.

**Figure 4:**
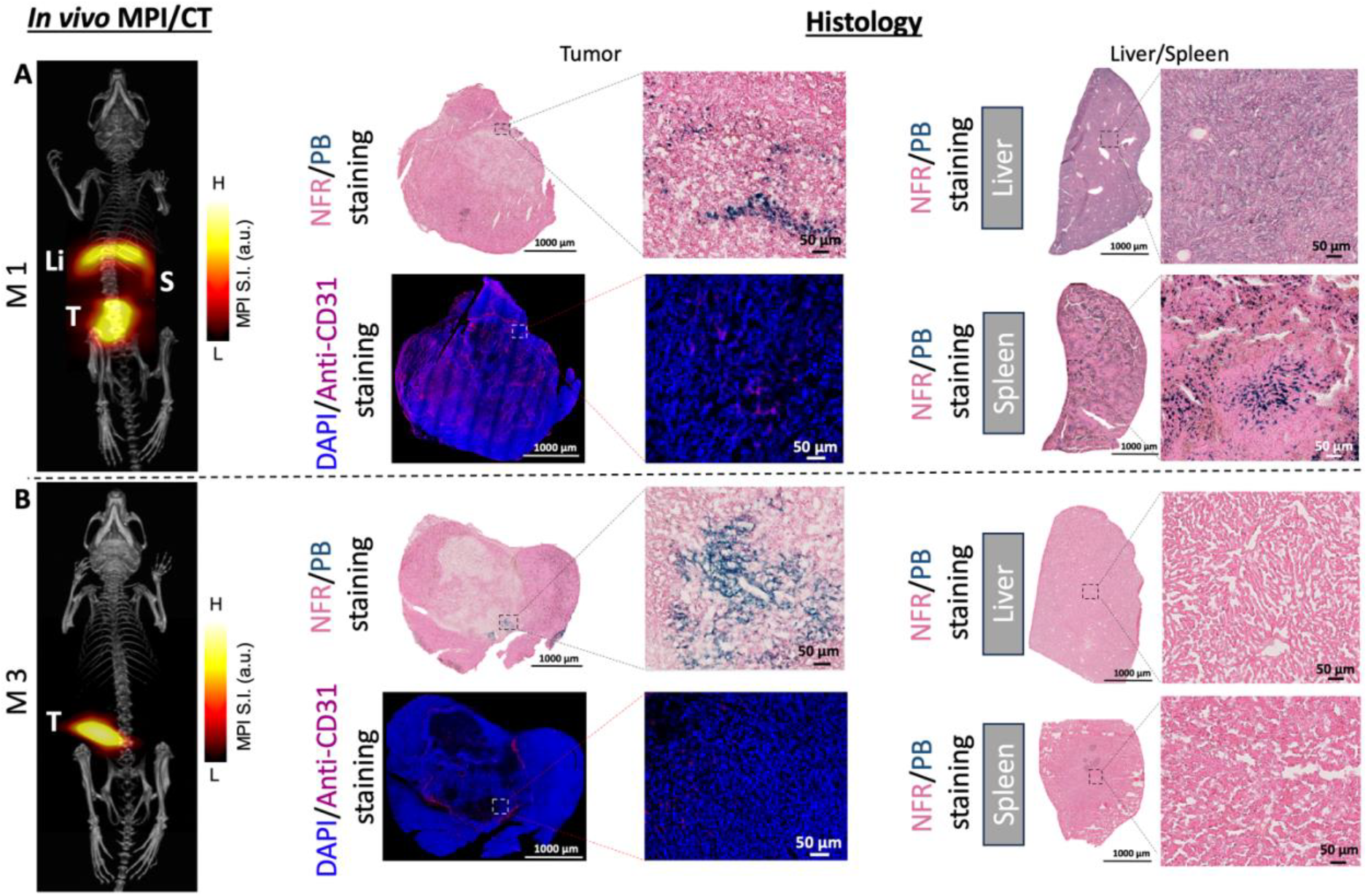
*In vivo* MPI/CT imaging and histological validation of MNP retention and systemic redistribution eight days post i.t. injection. Left panels show representative co-registered *in vivo* MPI/CT images acquired at the terminal time point, demonstrating MNP localization within tumors (T) in M1 (**A**) and M3 (**B**), with heterogeneous redistribution to the liver (Li) and spleen (S). Middle panels show histological analysis of excised tumors. Prussian blue (PB)/nuclear fast red (NFR) staining confirms the presence of MNP in tumor or organ tissues. Anti-CD31 immunostaining identifies tumor vasculature, enabling assessment of MNP distribution relative to blood vessels showing heterogeneous i.t. accumulation and retention. Right panels display PB staining of liver and spleen, demonstrating high inter-subject variability in MNP leakage and retention.

Quantitative image analysis further demonstrated substantial inter-subject differences in MNP retention (**Table 1**). The PB-positive coverage measured in M1 compared with M3 indicated a higher local PB-positive area within the analyzed section of M3. Immunofluorescence staining for CD31 revealed differences in vascular density between tumors that may explain this variability. CD31-positive coverage in M1 compared with M3, showed a higher vascular fraction in M1. This localized trend appeared inversely associated with PB-positive area within the analyzed sections. These observations suggest that vascular accessibility may contribute to MNP leakage from the tumor, with increased vascularity facilitating redistribution into systemic circulation during or shortly after injection.^17, 18^

**Table 1.** Quantitative histological analysis of i.t. MNP retention and vascular density.

| Staining type | Subject | Positive staining area (%) | Relative ratio (M1/M3) |
| --- | --- | --- | --- |
| Prussian blue | M1 | 0.88% | 0.25× |
|  | M3 | 3.44% |  |
| Anti-CD31 | M1 | 17.00% | 3.78× |
|  | M3 | 4.50% |  |

While phagocytic uptake of MNPs by tumor-associated macrophages is well established and its retention can be strongly influenced by interactions with immune and stromal cells rather than tumor cells themselves,^36, 37^ the early hepatic signal observed in M1 as early as 15 min following i.t. injection (**Fig. 2**) is consistent with rapid uptake of the injected material by liver Kupffer cells.^18^ This early liver signal may have resulted from injection-associated entry into the systemic circulation, including partial delivery into or near a blood vessel, leakage through highly vascularized tumor regions, or pressure-driven redistribution during injection. While we cannot exclude i.t. MNP uptake by tumor macrophages with subsequent systemic redistribution, it is unlikely to have occurred at such early time points.

## CONCLUSIONS

We presented a quantitative longitudinal MPI approach for monitoring the fate of locally injected MNPs following i.t. delivery. Serial whole-body MPI enabled noninvasive tracking of MNP-associated signal over time and revealed dynamic changes in i.t. retention and systemic redistribution patterns across animals. *Ex vivo* MPI and histological analysis provided complementary information regarding organ-level signal distribution and localized NP deposition. The ability of MPI to directly measure NP-associated signal over time could be valuable for future studies investigating NP delivery heterogeneity, treatment-associated redistribution, and imaging-guided optimization of i.t. nanotherapeutic strategies.

## Acknowledgments

We thank Dr. Marzieh Salimi for experimental assistance with the tumor model.

## Author contributions

Conceptualization: JWMB, RI

Methodology: ASZ, AI, JG, PK, JWMB

Investigation: ASZ, AI, JG, PK

Visualization: ASZ, AI, JG, PK

Funding acquisition: RI, JWMB

Project administration: JWMB

Supervision: JWMB

Writing – original draft: ASZ, JWMB

Writing – review & editing: ASZ, AI, JG, PK, RI, JWMB

## Data and materials availability

All data needed to evaluate and reproduce the conclusions of this work are present in the paper. This study did not generate new materials. All correspondence should be sent to J.W.M.B.

